## Supplementary Information for "Snowprint: a predictive tool for genetic biosensor discovery"

Supplementary Figures 1 - 7

Supplementary Tables 1 - 4

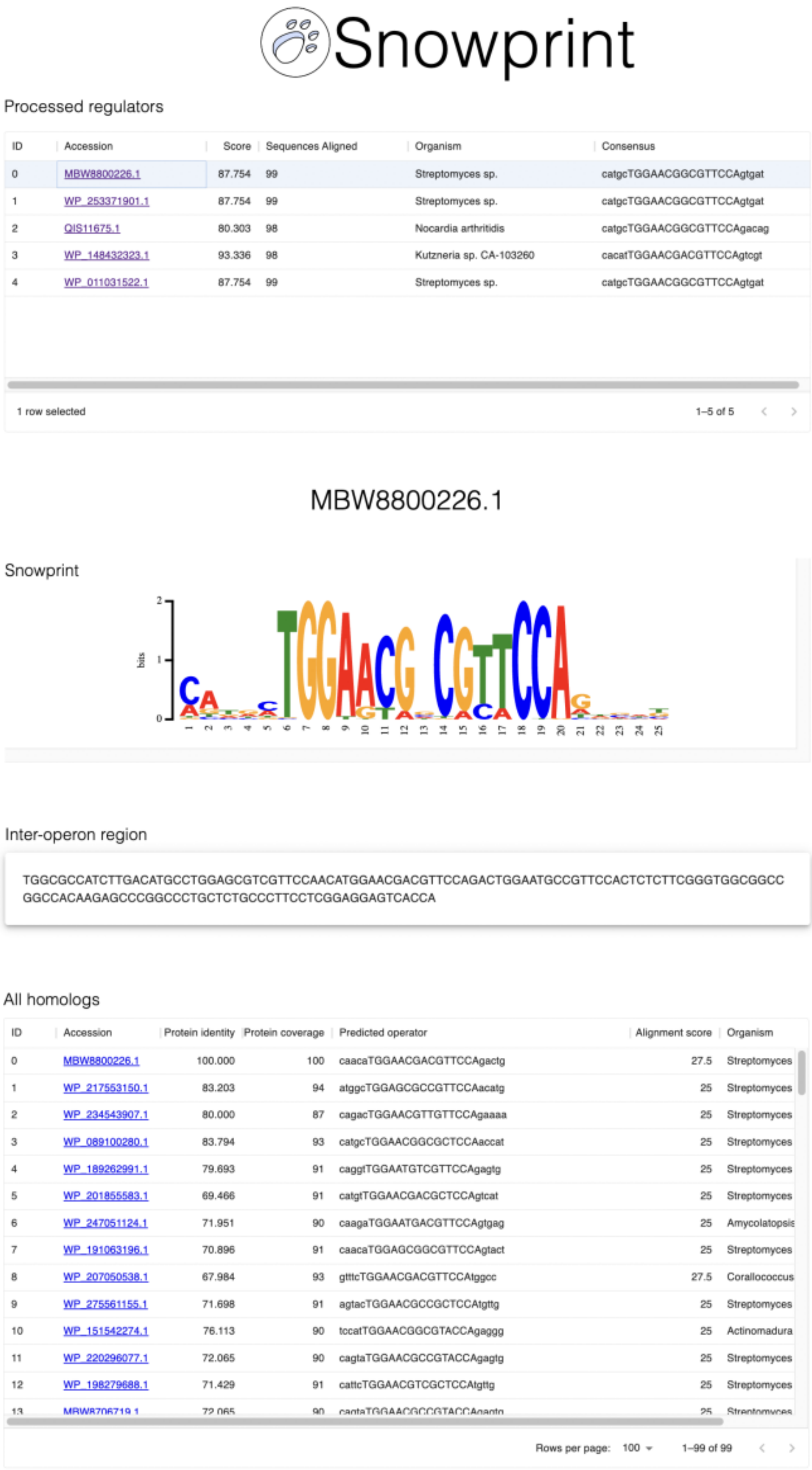


**Supplementary Figure 1**. React-based local web page displaying results from a Snowprint query.

Users can browse among predictions made for regulators run locally and view the (1) consensus prediction logo, (2) Inter-operon region containing the predicted operator, and (3) homologous predicted operators used to create the consensus prediction displayed in (1).

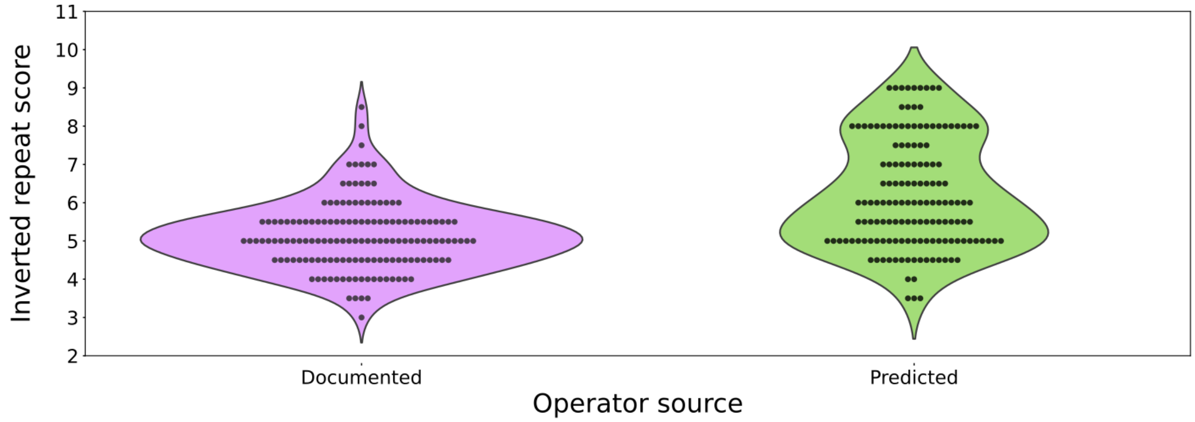


**Supplementary Figure 2**. Inverted repeat scores of operators within the benchmarking dataset.
Documented operators, shown in purple, represent the operator sequences extracted from literature (see **Supplementary Data 1**), while predicted operators, shown in green, represent operator sequences predicted by Snowprint using the associated regulator RefSeq accession ID.

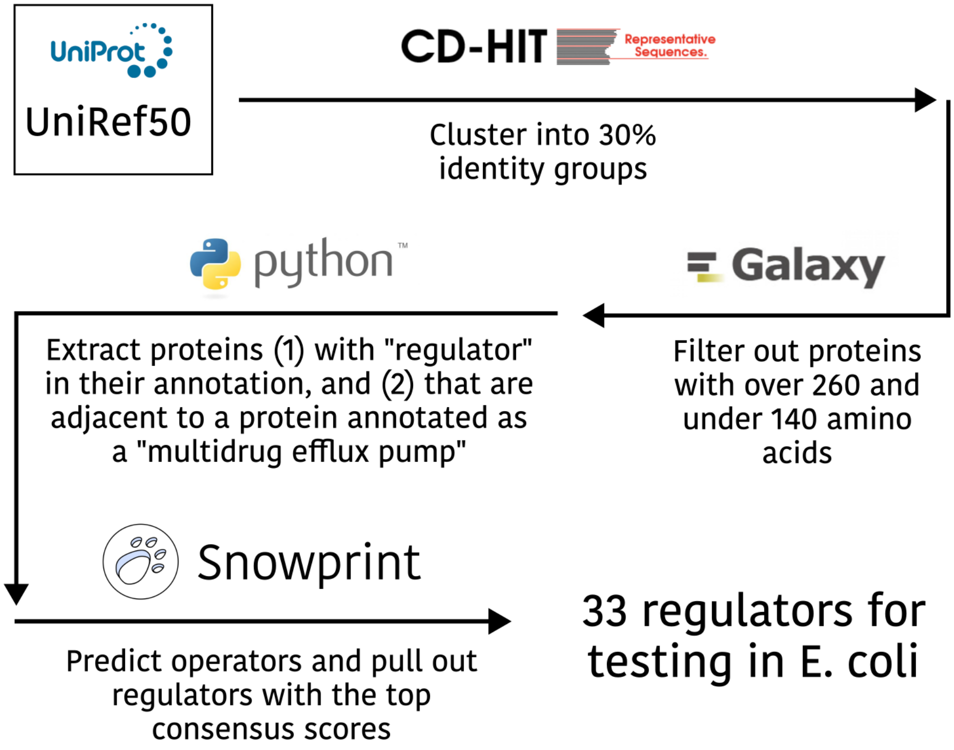


**Supplementary Figure 3**. Generalist regulator curation workflow.
TetR-family regulators were fetched from the UniRef50 sequence database, which were then clustered into 30% similarity groups using the CD-HIT software and filtered to contain only protein sequences shorter than 260 amino acids and longer than 140 amino acids using the Galaxy Suite software. Custom python scripts were then used with the Entrez API to filter proteins that contain “regulator” in their annotation and that are adjacent to proteins annotated as “multidrug efflux pumps”. The resulting regulators were then used as inputs to generate operator predictions with Snowprint, and the 33 regulators with the top consensus scores were selected for functional testing within *E. coli* cells.

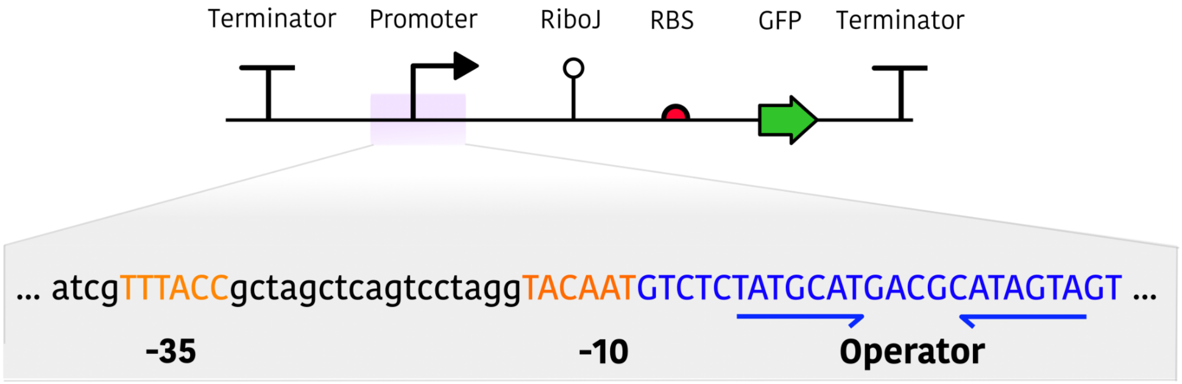


**Supplementary Figure 4**. Promoter design for domesticated regulators

Regulator-controlled promoters were designed by placing the predicted operator sequence immediately downstream from the -10 box of an E. coli sigma-70 promoter. Arrows indicate inverted repeat sequences, showing that five base pairs outside of the inverted repeat region were included in the promoter. The broader context of the reporter transcriptional unit is shown above the promoter sequence.

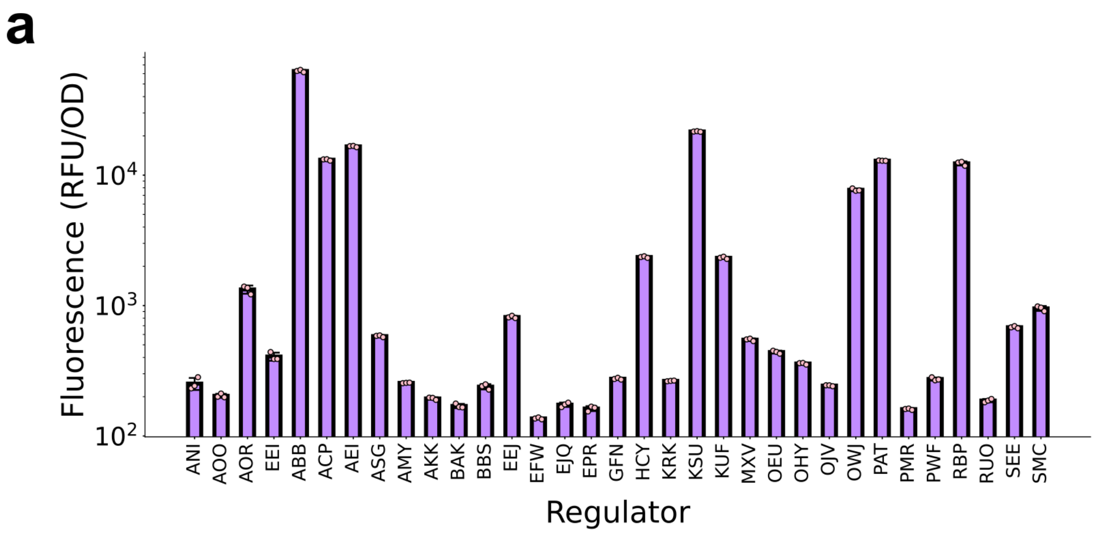


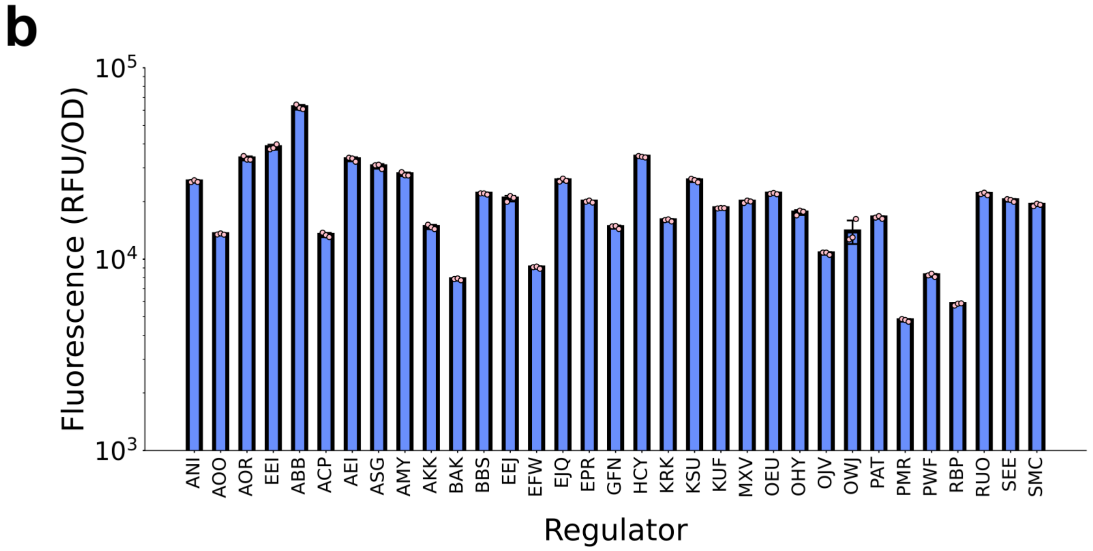


**Supplementary Figure 5**. Synthetic promoter output with and without domesticated regulator expression. Fluorescence of *E. coli* cells containing a reporter plasmid expressing GFP from a promoter bearing a Snowprint-predicted operator (see Figure 3, Supplementary Figure 4) and a regulator plasmid expressing a transcription factor from a constitutive plasmid. In (a) a regulator plasmid expresses the transcription factor predicted to bind to the Snowprint-predicted operator. In (b) the regulator plasmid expresses the CamR transcription factor. Assays were performed in biological triplicate. Individual data points are shown in pink. Error bars represent the standard deviation +/- the mean.

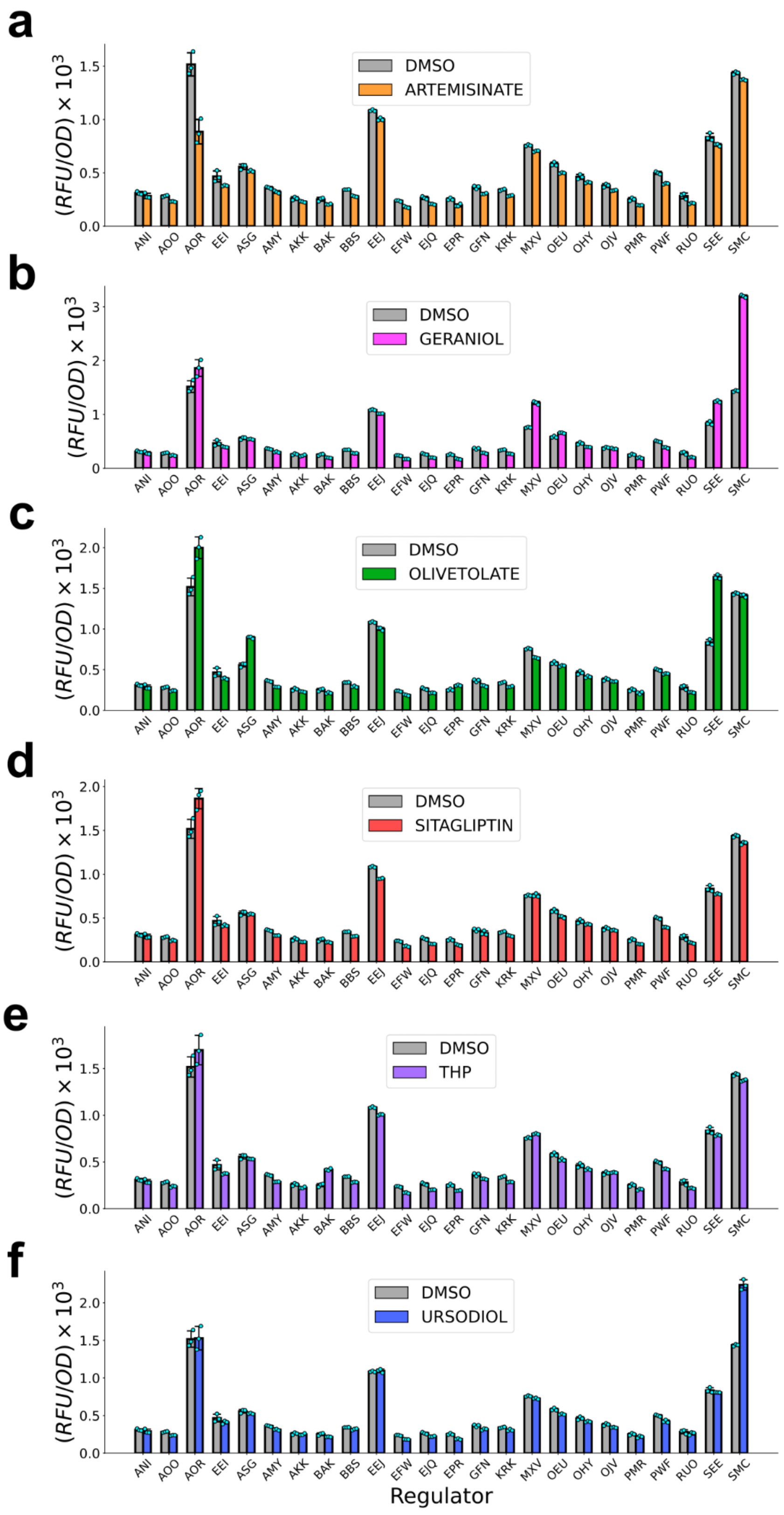


**Supplementary Figure 6**. Response of 24 generalist regulators to biomanufacturing-relevant ligands. The final concentration for induction of each ligand was 100 uM.   Assays were performed in biological triplicate. Individual data points are shown in cyan. Error bars represent the standard deviation +/- the mean.

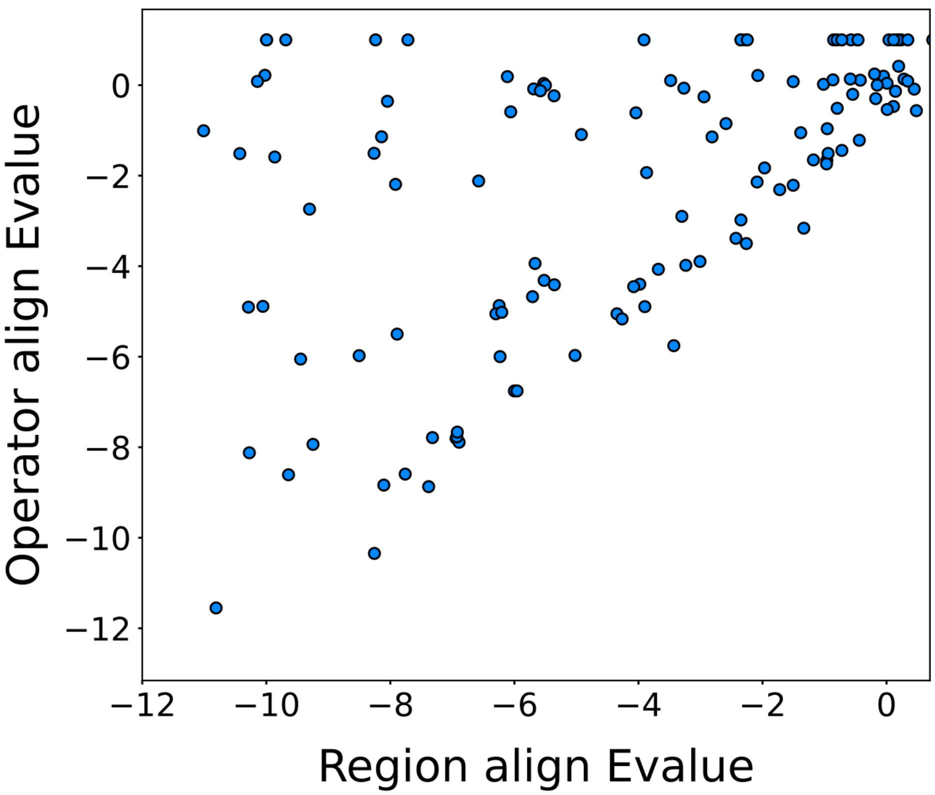


**Supplementary Figure 7**. Comparison of known operator alignment to the predicted operator vs the inter-operon region. Single data points represent the E-value similarity score of a known operator for its corresponding Snowprint predicted operator and the inter-operon region within the host organism's genome predicted to contain the operator.

<table>
<tr><th>Protein ID</th><th>E-value</th><th>Alignment</th></tr>
<tr><td>WP_001224188.1</td><td>$3.7*10^{-13}$</td><td>Documented 1 gttttataATAAACGGAgagttaTCCGTTTGTcaa 35<br>||||||||||||.||||||||||||||||<br>Predicted 1 -----ataatAAACGGATAGTTATCCGTTTgtcaa 30</td></tr>
<tr><td>AEM66515.1</td><td>0.0012</td><td>Documented 1 TAGCCACGTCTGGACAAAGTGAGAGATCGTGTCTAGACAACGCCACGGTT 50<br>||||||||||||||||||||<br>Predicted 1 -----------------------gatcgTGTCTAGACAacgcc------ 20</td></tr>
<tr><td>NP_415533.1</td><td>0.0076</td><td>Documented 1 TGTAAATTTTGACCATTTGGTCCACTTTTTTCT 33<br>.|||.||||.|..|..|||.<br>Predicted 1 -TTAACTTTTTAAAAACTGGC------------ 20</td></tr>
<tr><td>WP_011060270.1</td><td>0.0064</td><td>Documented 1 -----TCAAACAAGTGTTTGTCAGG--- 20<br>|.||||...||||  |||..<br>Predicted 1 aagccTGAAACGTATGTT--TCAaacaa 26</td></tr>
<tr><td>WP_005058758.1</td><td>0.0072</td><td>Documented 1 ---------CTTAACGCATGGCATGTGCGTTATG 25<br>|.|.|||||.|.|||.<br>Predicted 1 ggtagCGTGCGTTACGCACGtcata--------- 25</td></tr>
<tr><td>CAY46636.1</td><td>0.0049</td><td>Documented 1 ---GCT-----TGTATGTACAAGT-- 16<br>.||      |||||.|||||.|<br>Predicted 1 acgtcTGTATATGTATATACAaataa 26</td></tr>
<tr><td>NP_391277.1</td><td>0.0061</td><td>Documented 1 ---AAATTTGTCCGTATACATTTT-- 21<br>.||||.||.||||.|.||.|.<br>Predicted 1 atttaATTAGTACGTACAAATataga 26</td></tr>
<tr><td>NP_227848.1</td><td>0.0018</td><td>Documented 1 --AATTTCTTTCTGAG-GAAGATAGA 23<br>|..||||....|.| |||||..|<br>Predicted 1 agacaTTCTCAAAGTGAGAAgaatg- 25</td></tr>
<tr><td>WP_003084488.1</td><td>1.0</td><td>Documented 1 tttcTAG--AGTAaaTACTCTAaagt-- 24<br>||..|  .|......|.|...|.|<br>Predicted 1 --tcacgGCGGGGCGCGCCCCGCactgt 26</td></tr>
</table>

**Supplementary Table 1.** Global alignment of documented and predicted operators.
The first and last alignments display excellent and poor scores for comparison, respectively. All other alignments have E-values between 0.01 and 0.001. The pairwise EMBOSS needle algorithm was used to create alignments with default parameters. Documented operators represent sequences experimentally validated to bind the corresponding protein regulator. Refer to Supplementary Data 1 for more information on sequence metrics and sources.

| Genbank ID | Number of aligned sequences | Consensus Score | Predicted operator |
|---|---|---|---|
| ANI80042.1 | 7 | 94.99 | gttcgAAAATAAACAATCTTGTTTCTTTTattca |
| AOO80349 | 12 | 89.099 | cctgcTAAACTAAACGGTTCAGTTTAtttcg |
| AOR61801.1 | 10 | 62.821 | tatacCCATAATTAATGACCCAGAAGTCATTAATTATGGtgtcc |
| EEI22781.1 | 38 | 95.314 | ctcatAGTGACACTGTGTCACTcgatt |
| ABB11310.1 | 24 | 85.375 | cgattTGCGTGCCCCGGGTGCCGGGGCACGCAtcctg |
| ACP22327 | 15 | 75.794 | cttatTAGATTTCAATTGACATCTAtatat |
| AEI82798.1 | 38 | 71.203 | cttTGCGGTCCcGGACCGCAtga |
| ASG24684.1 | 9 | 100 | ggactGTACTGTATGATACAGTACgtaat |
| AMY71450 | 11 | 82.012 | caatcACTGTACCGTCCAGACGGTAAAGTcaacc |
| AKK09602.1 | 24 | 68.741 | acataTATTTCATCAACCGATGAAATAcggag |
| BAK71752.1 | 36 | 89.688 | tttatAATACATACGTATGTATTtaata |
| BBS35964.1 | 8 | 71.611 | agataAGCTAAACTTGAATAAAGATTCAAGTTTAGCTacttt |
| EEJ50829.1 | 21 | 76.158 | ggtgtGGTGACAAGTTGTCACCggttg |
| EFW06302.1 | 70 | 72.686 | cctttTTTAAATGAATATTTATTCATTCAAAttata |
| EJQ60916.1 | 99 | 92.352 | attgtCTAACAGTGTTAGtttga |
| EPR37357.1 | 9 | 90.361 | tccagAACTGACCGGTCAGTTtttga |
| GFN32285.1 | 9 | 85.012 | acaatATGAATGAATCATTCATttcga |
| HCY59852 | 24 | 81.386 | aaACCTAGTAAGTTACTAGGTga |
| KRK48889 | 12 | 96.543 | aacggTTGTAAATTTAAATTACAAagagt |
| KSU66562.1 | 34 | 56.409 | aaacaGGTGAGCGCCCGAGCATTATACTCGGGCGCTCACCcgttt |
| KUF06535.1 | 18 | 86.464 | agaatATACCCATCGGTATctagg |
| MXV43606.1 | 21 | 82.634 | ttagcGTGGAAACCGATTAGTTTCCACccccct |
| OEU66612 | 6 | 84.907 | aacacTTGTTTAAAACAAtcgtt |
| OHV36883.1 | 13 | 87.706 | ctactACCGTCCGTCCGGACGGTcggtc |
| OJV33711 | 28 | 84.969 | gcaaaACGTGAGTGAACGATCACTCACGTactag |
| OWJ67207.1 | 100 | 45.685 | cccgtCCGCCCCCGATTTCTCGGGGTCGGcgctg |
| PAT04833.1 | 12 | 95.26 | cgcccCAAGGTATTACCTTGcacac |
| PMR76500 | 16 | 96.12 | tgttgTTAACGCCGTTAAtatag |
| PWF24669 | 15 | 62.766 | ctctaAAATTAGGTCGGTTAACCAACCTAATTTtatgc |
| RBP44292.1 | 3 | 86.607 | tttacAGGTATGCAAGCAAGTACTTACTTGCATACCTgttgc |
| RUO21779.1 | 11 | 71.091 | gaatcTCGGACTGACTAGTCCGTCCGAtggtt |
| SEE04737 | 41 | 76.365 | attgaATTGCACCAGATAGTGCAATaggag |
| SMC09139 | 8 | 90.222 | ccTTGTGAATgagATTCACAAaa |

**Supplementary Table 2.** Snowprint predictions for mined regulators.
Results and metadata associated with Snowprint predictions of TetR-family transcriptional regulators targeted for testing within *E. coli*. Number of aligned sequences represents the number of homologous operator sequences used to generate the consensus. See **Methods** for consensus score calculation.

TGCTTCATTCCTCGGTACCAAATTCCAGAAAAGAGGGGAGCGGAAACCGCTCCCCTTTTTCGTTTTGGTCCCTCGC
TGCAGTGTCGTAAAAACCAGTTGGGGAGACGCTGTGAATCCCGGGCAAGCAAAATCGCTGGAAGTTCCTATACTTTC
TAGAGAATAGGAACTTCTTTCTAAATACATTCAAATATGTATCCGCTCATGAGACAATAACCCTGATAAATGCTTCAA
TAATATTGAAAAAGGAAGAGTATGAGCCATATTCAACGGGAAACGTCTTGCTCCAGGCCGCGATTAAATTCCAACAT
GGATGCTGATTTATATGGGTATAAATGGGCTCGCGATAATGTCGGGCAATCAGGTGCGACAATCTATCGATTGTATGG
GAAGCCCGATGCGCCAGAGTTGTTTCTGAAACATGGCAAAGGTAGCGTTGCCAATGATGTTACAGATGAGATGGTCA
GACTAAACTGGCTGACGGAATTTATGCCTCTTCCGACCATCAAGCATTTTATCCGTACTCCTGATGATGCATGGTTAC
TCACCACTGCGATCCCCGGGAAAACAGCATTCCAGGTATTAGAAGAATATCCTGATTCAGGTGAAAATATTGTTGATG
CGCTGGCAGTGTTCCTGCGCCGGTTGCATTCGATTCCTGTTTGTAATTGTCCTTTTAACAGCGATCGCGTATTTCGCC
TCGCTCAGGCGCAATCACGAATGAATAACGGTTTGGTTGATGCGAGTGATTTTGATGACGAGCGTAATGGCTGGCCT
GTTGAACAAGTCTGGAAAGAAATGCATAAGCTTTTGCCATTCTCACCGGATTCAGTCGTCACTCATGGTGATTTCTC
ACTTGATAACCTTATTTTTGACGAGGGGAAATTAATAGGTTGTATTGATGTTGGACGAGTCGGAATCGCAGACCGATA
CCAGGATCTTGCCATCCTATGGAACTGCCTCGGTGAGTTTTCTCCTTCATTACAGAAACGGCTTTTTCAAAAATATGG
TATTGATAATCCTGATATGAATAAATTGCAGTTTCATTTGATGCTCGATGAGTTTTTCTAAGTTGTGATGGCGGTAGG
AATGTAATCGTTAATCCGCAAATAACGTAAAAACCCGCTTCGGCGGGTTTTTTTATGGGGGGAGTTTAGGGAAAGAG
CATTTGTCATCCCGTTGAATATGGCTCCCTTAACGTGAGGAAGTTCCTATACTTTCTAGAGAATAGGAACTTCTACAG
ATGGACTTGGGTTGGCGGTTTCAGGAGTAGGTGCTTCCTCGCTCACTGACTCGCTGCACGAGGCAGACCTCAGCGCT
AGCGGAGTGTATACTGGCTTACTATGTTGGCACTGATGAGGGTGTCAGTGAAGTGCTTCATGTGGCAGGAGAAAAAA
GGCTGCACCGGTGCGTCAGCAGAATATGTGATACAGGATATATTCCGCTTCCTCGCTCACTGACTCGCTACGCTCGGTC
GTTCGACTGCGGCGAGCGGAAATGGCTTACGAACGGGGCGGAGATTTCCTGGAAGATGCCAGGAAGATACTTAACAG
GGAAGTGAGAGGGCCGCGGCAAAGCCGTTTTTCCATAGGCTCCGCCCCCCTGACAAGCATCACGAAATCTGACGCTCA
AATCAGTGGTGGCGAAACCCGACAGGACTATAAAGATACCAGGCGTTTCCCCCTGGCGGCTCCCTCGTGCGCTCTCCT
GTTCCTGCCTTTCGGTTTACCGGTGTCATTCCGCTGTTATGGCCGCGTTTGTCTCATTCCACGCCTGACACTCAGTTC
CGGGTAGGCAGTTCGCTCCAAGCTGGACTGTATGCACGAACCCCCCGTTCAGTCCGACCGCTGCGCCTTATCCGGTAA
CTATCGTCTTGAGTCCAACCCGGAAAGACATGCAAAAGCACCACTGGCAGCAGCCACTGGTAATTGATTTAGAGGAG
TTAGTCTTGAAGTCATGCGCCGGTTAAGGCTAAACTGAAAGGACAAGTTTTGGTGACTGCGCTCCTCCAAGCCAGTT
ACCTCGGTTCAAAGAGTTGGTAGCTCAGAGAACCTTCGAAAAACCGCCCTGCAAGGCGGTTTTTTCGTTTTCAGAGC
AAGAGATTACGCGCAGACCAAAACGATCTCAAGAAGATCATCTTATTAAGGGGTCTGACGCTCAGTGGAACGAAACC
CTGTTGGTCAGGTTTTCGGGAGGTGTGCGTCTCGCATCCGGAAAGGTGTGATAGGTAGCGCAGCAAATAAAACGAAA
GGCTCAGTCGAAAGACTGGGCCTTTCGTTTTATCTGTTGTTTGTCGGTGAACGCTCTCCTCAACGAAAATATATTTTT
CAAAAGTATCGTTTACCGCTAGCTCAGTCCTAGGTACAATNNNNNNNNNNNNNNNNNTGGCAGCTGTCACCGGAT
GTGCTTTCCGGTCTGATGAGTCCGTGAGGACGAAACAGCCTCTACAAATAATTTTGTTTAAGGGCCCAAGTTCACTTA
AAAAGGAGATCAACAATGAAAGCAATTTTCGTACTGAAACATCTTAATCATGCGGTAAGGAGTTAAATATGTCGAAG
GGAGAGGAATTGTTTACTGGAGTCGTCCCAATTCTTGTTGAGTTGGACGGAGATGTCAATGGTCATAAGTTTAGCGT
ATCCGGTGAGGGTGAGGGAGATGCTACTTACGGAAAATTAACATTAAAATTCATTTGTACCACAGGCAAACTTCCTGT
TCCTTGGCCAACTCTTGTGACGACTTTTGGCTACGGTGTGCAGTGTTTCGCTCGTTATCCTGACCACATGAAGCAACA
TGATTTCTTCAAGTCTGCGATGCCTGAAGGATATGTTCAAGAACGTACCATCTTCTTCAAAGATGATGGTAACTATAA
GACTCGTGCAGAGGTAAAATTCGAGGGAGATACGTTGGTAAATCGCATTGAGCTGAAGGGAATCGATTTCAAGGAAG
ATGGAAACATTCTGGGACACAAGCTGGAGTACAATTACAATAGCCATAACGTCTATATCATGGCAGATAAGCAAAAAA
ACGGAATCAAGGTTAACTTCAAGATTCGCCATAATATTGAGGACGGCTCTGTGCAATTGGCGGATCATTATCAACAGA
ACACCCCGATTGGAGATGGTCCCGTGCTGCTGCCAGATAATCACTATTTGTCTACACAATCGGCCCTTTCCAAAGATC
CGAACGAAAAGCGCGATCATATGGTACTTTTAGAGTTCGTAACTGCCGCTGGAATCACCCACGGCATGGATGAGTTGT
ATAAGTAATAATCC

p15A origin
Kanamycin resistance gene
GFP
Promoter
Predicted operator
RiboJ
Terminator

**Supplementary Table 3**. Full plasmid sequence of the reporter vector

AAAACTCACGTTAAGGCCCTCTCCAAGACCGAGCCATCAACAAAGCGTCTCGCTGAGGTTTCATGGAGCCTCTGGTTCATCTCCGG
CAATTAAAAAAGCGGCTAACCACGCCGCTTTTTTTACGTCTGCAGGAACGGGCTGTCGACCTTTGAAAAGTTCGTTTACCGCTAGC
TCAGTCCTAGGTACAATNNNNNNNNNNNNNNNNNNTGGCAGCTGTCACCGGATGTGCTTTCCGGTCTGATGAGTCCGTGAGGAC
GAAACAGCCTCTACAAATAATTTTGTTTAAGGGCCCAAGTTCACTTAAAAAGGAGATCAACAATGAAAGCAATTTTCGTACTGAAA
CATCTTAATCATGCGGAGGCAGCTTAAATATGTCTACTAAGAACAAGATTGACAAGAACTATCTGATCAACATCATCGAGGAGATCC
TGCTGAACGATGGAATTAGTGGCTTGAGCATTCGTAAAGTAGCCACTAAAGCTAATATCTCAATCGGTGGTGTTCAGTATATCTTCG
GCAATAAGGAAGGGATGATCAAAGCGGTCCTTGAAAAAAACGAAGAGGATTACAACCGCCAGATTAAAATTCTCCTGAAGGATGA

CAATAGCAAATACTCCCAGTTGAAGGCGCATATCGAGTATATTTCGAAGCACAACGATAATGAGGAGTTCGACAAAATCTCGAAGA
TTATTACGATTCTGCTCCAAGAGAAATTCGTGTTCGAAGGTCTGCAAGATTGGTACAGTACGTCTTTAAATAGCATTGACACCAATA
CCGATGAGGGCAAAAAATTGCGGCTGGCATTTCTGTTCTCCGAAGCCATGTTTACACTGATGACCTTAAAGTATATCAACATTAGCC
CGAAGGAACAGAAGGAGATTTTCGAAGATCTGAAAACCTTTTTACTTTAATAATCCTATCGCCACTTTCAGCCAAAAAACTTAAGA
CCGCCGGTCTTGTCCACTACCTTGCAGTAATGCGGTGGACAGGATCGGCGGTTTTCTTTTCTCTTCTCAACACCCTTCGCGTCAACA
CTTTTCCGCCAAGGAGACGGTTGGTCAGGTTTTCGGGAGGTGTGGCTGGAAGTTCCTATACTTTCTAGAGAATAGGAACTTCTTTC
TAAATACATTCAAATATGTATCCGCTCATGAGACAATAACCCTGATAAATGCTTCAATAATATTGAAAAAGGAAGAGTATGAGTATT
CAACATTTCCGTGTCGCCCTTATTCCCTTTTTTGCGGCATTTTGCCTTCCTGTTTTTGCTCACCCAGAAACGCTGGTGAAAGTAAA
AGATGCTGAAGATCAGTTGGGTGCACGAGTGGGTTACATCGAACTGGATCTCAACAGCGGTAAGATCCTTGAGAGTTTTCGCCCCG
AAGAACGTTTTCCAATGATGAGCACTTTTAAAGTTCTGCTATGTGGCGCGGTATTATCCCGTGTTGACGCCGGGCAAGAGCAACTC
GGTCGCCGCATACACTATTCTCAGAATGACTTGGTTGAGTACTCACCAGTCACAGAAAAGCATCTTACGGATGGCATGACAGTAAG
AGAATTATGCAGTGCTGCCATAACCATGAGTGATAACACTGCGGCCAACTTACTTCTGACAACGATCGGAGGACCGAAGGAGCTAA
CCGCTTTTTTGCACAACATGGGGGATCATGTAACTCGCCTTGATCGTTGGGAACCGGAGCTGAATGAAGCCATACCAAACGACGAG
CGTGACACCACGATGCCTGCAGCAATGGCAACAACGTTGCGCAAACTATTAACTGGCGAACTACTTACTCTAGCTTCCCGGCAACA
ATTAATAGACTGGATGGAGGCGGATAAAGTTGCAGGACCACTTCTGCGCTCGGCCCTTCCGGCTGGCTGGTTTATTGCTGATAAATC
TGGAGCCGGTGAGCGTGGATCGCGCGGTATCATTGCAGCACTGGGGCCAGATGGTAAGCCCTCCCGTATCGTAGTTATCTACACGAC
GGGGAGTCAGGCAACTATGGATGAACGAAATAGACAGATCGCTGAGATAGGTGCCTCACTGATTAAGCATTGGTAAGTTGTGATGG
CGGTAGGAATGTAATCGTTAATCCGCAAATAACGTAAAAACCCGCTTCGGCGGGTTTTTTTATGGGGGGAGTTTAGGGAAAGAGCA
TTTGTCATCCCGTTGAATATGGCTCCCTTAACGTGAGGAAGTTCCTATACTTTCTAGAGAATAGGAACTTCTACAGATGGACTTGGG
TTGGCGGTTTCAGGAGTCTGCAAAACGTCTGCGACCTGAGCAACAACATGAATGGTCATCGGTTTCCGTGTTTCGTAAAGTCTGGA
AACGCGGAAGTCAGCGCCCTGCACCATTATGTTCCGGATCTGCATCGCAGGATGCTGCTGGCTACCCTGTGGAACACCTACATCTGT
ATTAACGAAGCGCTGGCATTGACCCTGAGTGATTTTTCTCTGGTCCCGCCGCATCCATACCGCCAGTTGTTTACCCTCACAACGTTC
CAGTAACCGGGCATGTTCATCATCAGTAACCCGTATCGTGAGCATCCTCTCTCGTTTCATCGGTATCATTACCCCCATGAACAGAAA
TCCCCCTTACACGGAGGCATCAGTGACCAAACAGGAAAAAACCGCCCTTAACATGGCCCGCTTTATCAGAAGCCAGACATTAACGC
TTCTGGAGAAACTCAACGAGCTGGACGCGGATGAACAGGCAGACATCTGTGAATCGCTTCACGACCACGCTGATGAGCTTTACCGC
AGCTGCCTCGCGCGTTTCGGTGATGACGGTGAAAACCTCTGACACATGCAGCTCCCGCAGACGGTCACAGCTTGTCTGTAAGCGGA
TGCCGGGAGCAGACAAGCCCGTCAGGGCGCGTCAGCGGGTGTTGGCGGGTGTCGGGGCGCAGCCATGACCCAGTCACGTAGCGATA
GCGGAGTGTATACTGGCTTAACTATGCGGCATCAGAGCAGATTGTACTGAGAGTGCACCGGTGTGAAATACCGCACAGATGCGTAAG
GAGAAAATACCGCATCAGGCGCTCTTCCGCTTCCTCGCTCACTGACTCGCTGCGCTCGGTCGTTCGGCTGCGGCGAGCGGTATCAGC
TCACTCAAAGGCGGTAATACGGTTATCCACAGAATCAGGGGATAACGCAGGAAAGAACATGTGAGCAAAAGGCCAGCAAAAGGCC
AGGAACCGTAAAAAGGCCGCGTTGCTGGCGTTTTTCCATAGGCTCCGCCCCCCTGACGAGCATCACAAAAATCGACGCTCAAGTCA
GAGGTGGCGAAACCCGACAGGACTATAAAGATACCAGGCGTTTCCCCCTGGAAGCTCCCTCGTGCGCTCTCCTGTTCCGACCCTGCC
GCTTACCGGATACCTGTCCGCCTTTCTCCCTTCGGGAAGCGTGGCGCTTTCTCATAGCTCACGCTGTAGGTATCTCAGTTCGGTGTA
GGTCGTTCGCTCCAAGCTGGGCTGTGTGCACGAACCCCCCGTTCAGCCCGACCGCTGCGCCTTATCCGGTAACTATCGTCTTGAGTC
CAACCCGGTAAGACACGACTTATCGCCACTGGCAGCAGCCACTGGTAACAGGATTAGCAGAGCGAGGTATGTAGGCGGTGCTACAG
AGTTCTTGAAGTGGTGGCCTAACTACGGCTACACTAGAAGGACAGTATTTGGTATCTGCGCTCTGCTGAAGCCAGTTACCTTCGGA
AAAAGAGTTGGTAGCTCTTGATCCGGCAAACAAACCACCGCTGGTAGCGGTGGTTTTTTTGTTTGCAAGCAGCAGATTACGCGCAG
AAAAAAAGGATCTCAAGAAGATCCTTTGATCTTTTCTACGGGGTCTGACGCTCAGTGGAACG

pBR322 + ROP origin | Ampicillin resistance gene
Regulator (BAK as example) | Promoter
Predicted operator | RiboJ | Terminator

**Supplementary Table 4**. Full plasmid sequence of the regulator expression vector
